# BIOLOGICAL SCIENCES: Neuroscience, cell biology MTSS1/Src family kinase Dysregulation Underlies Multiple Inherited Ataxias

**DOI:** 10.1101/338046

**Authors:** Alexander S. Brown, Pratap Meera, Banu Altindag, Ravi Chopra, Emma Perkins, Sharan Paul, Daniel R. Scoles, Eric Tarapore, Mandy Jackson, Vikram G. Shakkottai, Thomas S. Otis, Stefan M. Pulst, Scott X. Atwood, Anthony E. Oro

## Abstract

The genetically heterogeneous Spinocerebellar ataxias (SCAs) are caused by Purkinje neuron dysfunction and degeneration, but their underlying pathological mechanisms remain elusive. The Src family of non-receptor tyrosine kinases (SFK) are essential for nervous system homeostasis and are increasingly implicated in degenerative disease. Here we reveal that the SFK suppressor Missing-in-Metastasis (MTSS1) is a novel ataxia locus that links multiple SCAs. MTSS1 loss results in increased SFK activity, reduced Purkinje neuron arborization, and low basal firing rates, followed by cell death. Surprisingly, mouse models for SCA1, SCA2, and SCA5 show elevated SFK activity, with SCA1 and SCA2 displaying dramatically reduced MTSS1 protein levels through reduced gene expression and protein translation, respectively. Treatment of each SCA model with a clinically-approved Src inhibitor corrects Purkinje basal firing, and delays ataxia progression in MTSS1 mutants. Our results identify a common SCA therapeutic target and demonstrate a key role for MTSS1/SFK in Purkinje neuron survival and ataxia progression.

**Significance Statement:** The Src family of non-receptor tyrosine kinases (SFK) are essential for nervous system function, and may contribute to neurodegeneration. Spinocerebellar ataxias (SCAs) are neurodegenerative diseases where Purkinje neurons fire irregularly and degenerate leading to motor problems. We show that the SFK suppressor Missing-in-Metastasis (MTSS1) is a novel ataxia gene that links multiple SCAs. MTSS1 loss results in increased SFK activity, degenerating Purkinje neurons with low basal firing rates, and cell death. Surprisingly, mouse models for three different SCAs show elevated SFK activity, with SCA1 and SCA2 models displaying dramatically reduced MTSS1 protein levels. Treatment of each SCA model with SFK inhibitor corrects Purkinje basal firing, and delays ataxia progression in MTSS1 mutants. Our results identify a common link among disparate neurodegenerative diseases.

## Introduction

Neurons are non-dividing cells that depend on homeostatic regulation of protein, RNA, and metabolite turnover to permit dynamic synaptic connections that allow adaptation to changing environments. Loss of such mechanisms result in one of several hundred neurodegenerative disorders. Over 40 loci form the genetic basis for human Spinocerebellar Ataxia (SCA), a progressive motor disorder characterized by cerebellar atrophy and pervasive Purkinje neuron degeneration where patients experience poor coordination and balance, hand-eye coordination, dysarthria, and abnormal saccades.

One common phenotype prominent in multiple SCA animal models is the altered Purkinje neuron firing rates that precede motor impairment and cell death (1–3), with restoration of the normal firing rates reducing Purkinje neuron death and improving motor function (4, 5). Defects in many cell functions lead to SCA including effectors of transcription (6), translation (7), proteostasis (8, 9), calcium flux (10, 11), and cytoskeletal/membrane interactions (12, 13). An open question remains how the many SCA genes interact to control firing rates and cell survival, with a common target emerging as a ideal treatment for the genetically diverse etiologies.

One such therapeutic target is the class of Src family of non-receptor tyrosine kinases (SFKs). Several SFKs are expressed in the nervous system and have partially overlapping functions. While single mutants for *Src* or *Yes* kinase have no overt neuronal phenotype (14, 15), *Fyn* loss of function leads to increased Src activity and hippocampal learning and memory deficits (16, 17) Moreover, *Fyn;Src* double mutants rarely survive past birth and have severely disorganized cortical and cerebellar layers (15, 18). SFKs are post-translationally regulated through activating and inhibitory phosphorylation marks deposited by inhibitory kinases and removed by receptor tyrosine phosphatases in a context dependent manner (19, 20). SFK activation occurs rapidly in response to extracellular signals and in response to a variety of cellular stresses ranging from osmotic pressure (21) to tetanic stimulation (22). Additionally, SFKs are inappropriately active in disease states including Amyotrophic lateral sclerosis (23), Alzheimer disease (24), and Duchenne muscular dystrophy (25).

Missing-in-Metastasis (MTSS1) is one of the defining members of the I-BAR family of negative membrane curvature sensing proteins first identified as being deleted in metastatic bladder cancer (26). Although MTSS1 biochemically interacts with membranes and regulates the actin cytoskeleton (27), genetic studies reveal that MTSS1 functions in an evolutionarily conserved signaling cassette to antagonize Src kinase activity (28, 29). Disruption of the MTSS1/Src regulatory cassette results in endocytosis and polarization abnormalities demonstrated by defects in primary cilia dependent hedgehog signaling, and hair follicle epithelial migration(28). In tissues requiring MTSS1 function, levels of active MTSS1 are critical, as loss (26) or gain (30) of MTSS1 has been associated with metastasis and invasion. Regardless of the particular phenotype, an evolutionarily conserved property of MTSS1 mutants is that loss of MTSS1 function can be reversed through the removal or inhibition of Src kinases. This property was first demonstrated through double mutant analysis in the fly ovary, and subsequently in mammalian tissue culture using Src kinase inhibitors (28, 29). The availability of FDA-approved Src kinase inhibitors has led to the investigation of clinically relevant MTSS1 phenotypes with the hope of using SFK inhibitors to ameliorate them.

Although SFKs have been shown to regulate multiple classes of neurotransmitter receptors (31) they also function to control basic cytoskeletal components. Src regulates local actin polymerization (32) and endocytic receptor internalization (32–35). The actin cytoskeleton plays a critical role in cell signaling, proliferation, motility, and survival. Local, rather than global, actin dynamics control homeostatic synaptic signaling, and abnormalities in actin regulation underlie a diversity of psychiatric and neuronal diseases including Amyotrophic lateral sclerosis (36), Schizophrenia, Autism Spectrum Disorders (37), and motor dysfunction such as spinocerebellar ataxia (SCA) (38). A major challenge remains to understand how actin cytoskeletal regulation controls synaptic function and to develop improved therapeutics for these common and poorly-treated diseases.

Here we reveal that actin regulator and SFK antagonist *Mtss1* is a novel ataxia locus regulated by multiple SCA alleles that subsequently result in SFK hyper-activation. We show that clinically-available Src inhibitors correct Purkinje neuron firing rates and delay ataxia progression, demonstrating a novel and druggable role for the evolutionarily conserved MTSS1/SFK network in Purkinje neuron survival and ataxia progression.

## Results

### Mtss1 null mice display a progressive ataxia

Mtss1 functions in a many tissues, and previous mutant alleles disrupting 5’ exons resulted in mild lymphmagenesis (39), progressive kidney disease (40), and mild neurological phenotypes (41). Since Mtss1 has several possible internal promoters likely not affected by these mutant alleles (42) we generated a conditional mutant allele targeting the endophilin/Src interacting domain located in the final exon (*MIM^Ex15^*, **Fig 1A**) (28, 29). Germline deletion with HPRT-cre resulted in the loss of MTSS1 protein as detected by an antibody specific to the N-terminal IMD domain (30) (**Fig 1B**). The absence of detectable MTSS1 protein supports the idea that the *MIM^EX15^* is a true null allele of *Mtss1*.

**Figure 1.**
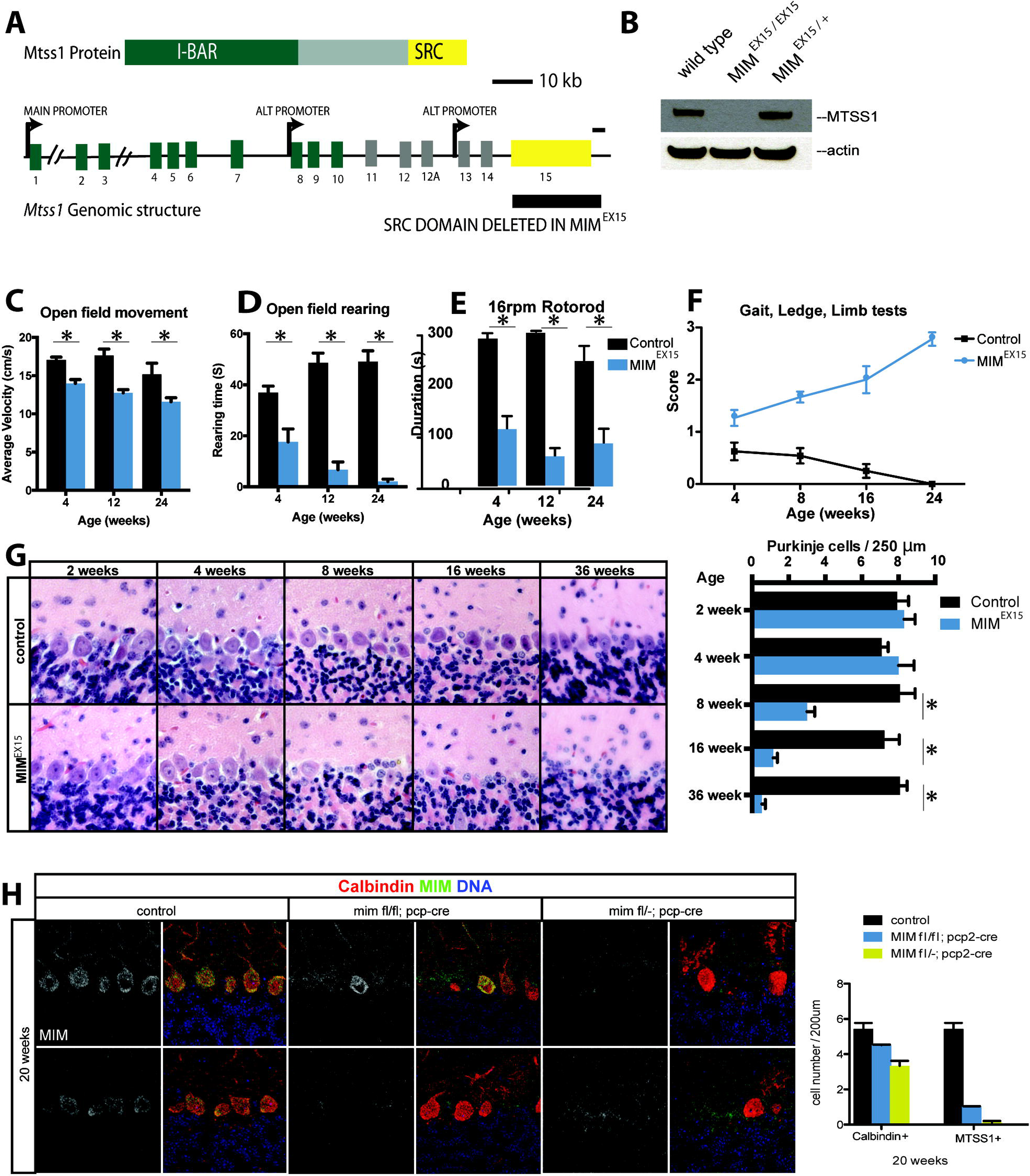
*MIM^EX15^* mutants develop progressive spinocerebellar ataxia. **A:** The structure of the *Mtss1* locus with alternative promoters and Src interacting domain deleted in *MIM^EX15^* mutants. **B**: Loss of MTSS1 protein in *MIM^EX15^* whole cerebellum lysate shown with N-terminal MTSS1 antibody. **C,D** *MIM^EX15^* show slower movement velocity and reduced rearing frequency in open field tests. **E:** Impaired rotarod learning in *MIM^EX15^* mutants, at the indicated ages, shown as reduced duration (time to fall).**F:** A composite test to measure spinocerebellar ataxia symptoms where increased score reflects reduced function showed an age dependent increase in severity in *MIM^EX15^* mutants. **G:** Age dependent loss of Purkinje neurons in *MIM^EX15^* mutants occurs after the onset of ataxia. **H:** At 20 weeks MIM^Loxp/-;^Pcp2-Cre and MIM^Loxp/Loxp;^Pcp2-Cre mutants show dramatic reduction in Purkinje neurons that stain with MTSS1. Many Purkinje neurons persist, as there is a less dramatic reduction in calbindin positive Purkinje cell number. Error bars, s.e.m.*p<0.05 students t-test. Error bars, s.e.m.

To our surprise, homozygous *MIM^EX15^* mutants lack cilia phenotypes and display a striking and progressive cerebellar ataxia. To better understand the nature of *MIM^EX15^* ataxia, we characterized *MIM^EX15^* mutants using an open field test to evaluate gross motor control. *MIM^EX15^* mutants had reduced velocity (**Fig 1C**) and rearing behavior (**Fig 1D**), consistent with overall movement defects. To uncouple possible motor and behavioral abnormalities we evaluated *MIM^EX15^* mutants with rotarod assay and observed coordination abnormalities in as early as 4 weeks of age (**Fig 1E**). Many spinocerebellar ataxias display progressive neurologic phenotypes. To determine whether *MIM^EX15^* animals showed progressive deterioration we employed a composite test measuring gait, grip strength and balance (43). We found *MIM^EX15^* animals performed consistently worse than controls, with severity increasing with age (**Fig 1F**). *MIM^EX15^* heterozygous animals displayed no overt phenotype.

Reduced Mtss1 levels are associated with a variety of cellular phenotypes including reduced presentation of receptors on the cell membrane (44), and altered Purkinje neuron morphology (41). To determine the basis of the motor abnormalities and to distinguish among these possibilities we performed histological analysis. At 4 weeks, *MIM^EX15^* mice are ataxic, yet their cerebella appeared grossly normal with intact granule, Purkinje neuron, and molecular layers. However, *MIM^EX15^* mutants displayed a profound and progressive loss of Purkinje neurons starting at 8 weeks of age. Whereas wild type cerebella contain approximately 8 Purkinje neurons in a 250 μm linear distance, 8-week old mice retained only 25% of wild type, and 24 week *MIM^EX15^* mutants contained only 5% of the total number of Purkinje neurons (**Fig 1G**).

While ataxia genes can act in many cell types to regulate Purkinje cell function, MTSS1 is highly expressed in Purkinje cells, suggesting it is required in these cells for normal Purkinje cell function and survival. To confirm the Purkinje neuron defects seen in *MIM^EX15^* animals are due to a cell autonomous requirement for *Mtss1*, we conditionally inactivated *Mtss1* using the Purkinje neuron specific L7-Cre (*MIM^cko^*) then compared Purkinje neuron morphology and loss to the global MIM^EX15^ mutant. *MIM^cko^* Purkinje neurons were mosaic for MTSS1 expression due to inefficient LoxP recombination. At 20 weeks *MIM^cko^* had reduced numbers of Purkinje neurons. In remaining Purkinje neurons, those lacking MTSS1 protein displayed thickened dendritic branches and reduced arbor volume, while neighboring Purkinje neurons with MTSS1 protein appeared normal (**Fig 1H**). We conclude that *Mtss1* acts cell autonomously in Purkinje neurons to maintain dendritic structure, with loss of MTSS1 resulting in abnormalities and eventual cell death.

### Mtss1 mutants have autophagocytic neurodegenerative disease

An emergent mechanism of cell loss during neurodegeneration is aberrant macroautophagy. Autophagy is essential for Purkinje neuron survival, as loss of autophagy (45, 46) results in cell death. Increased levels of early autophagy markers have been described in multiple neurodegenerative diseases including Huntington’s disease (47), Alzheimer disease (48), and SCA3 (49). *MIM^EX15^* mutants fit this pattern of disease as we observed multiple signs of autophagy. As early as 4 weeks, we observed increased Complex V/ATP synthase staining indicative of fused mitochondria as well as dramatically reduced staining for the Golgi body marker Giantin (**Figs 2A, S1**). By 8 weeks of age we could detect marked increased early autophagic marker LC3 and ubiquitinated proteins by immunofluorescence (**Figs 2B, S1**). We confirmed our immunofluorescence using electron microscopy of 8-week mutant cerebella. We observed Purkinje neurons with notched nuclei (**Fig 2C**), a phenotype associated with autophagy, rather than the shrunken, condensed nuclei associated with apoptosis. We also observed frequent cases of autophagic vacuoles and swollen mitochondria (**Fig 2D**), as well as late stage autophagic vacuoles filled with electron dense material (**Fig 2E**). Consistent with an autophagocytic cell death mechanism, we failed to see increased DNA breaks in *MIM^EX15^* Purkinje neurons with TUNEL stain (**Fig 2F**). In addition to autophagocytic neurodegeneration, *MIM^EX15^* animals displayed increased GFAP positive glial infiltration (**Fig 2G**) consistent with reactive astroglyosis.

**Figure 2.**
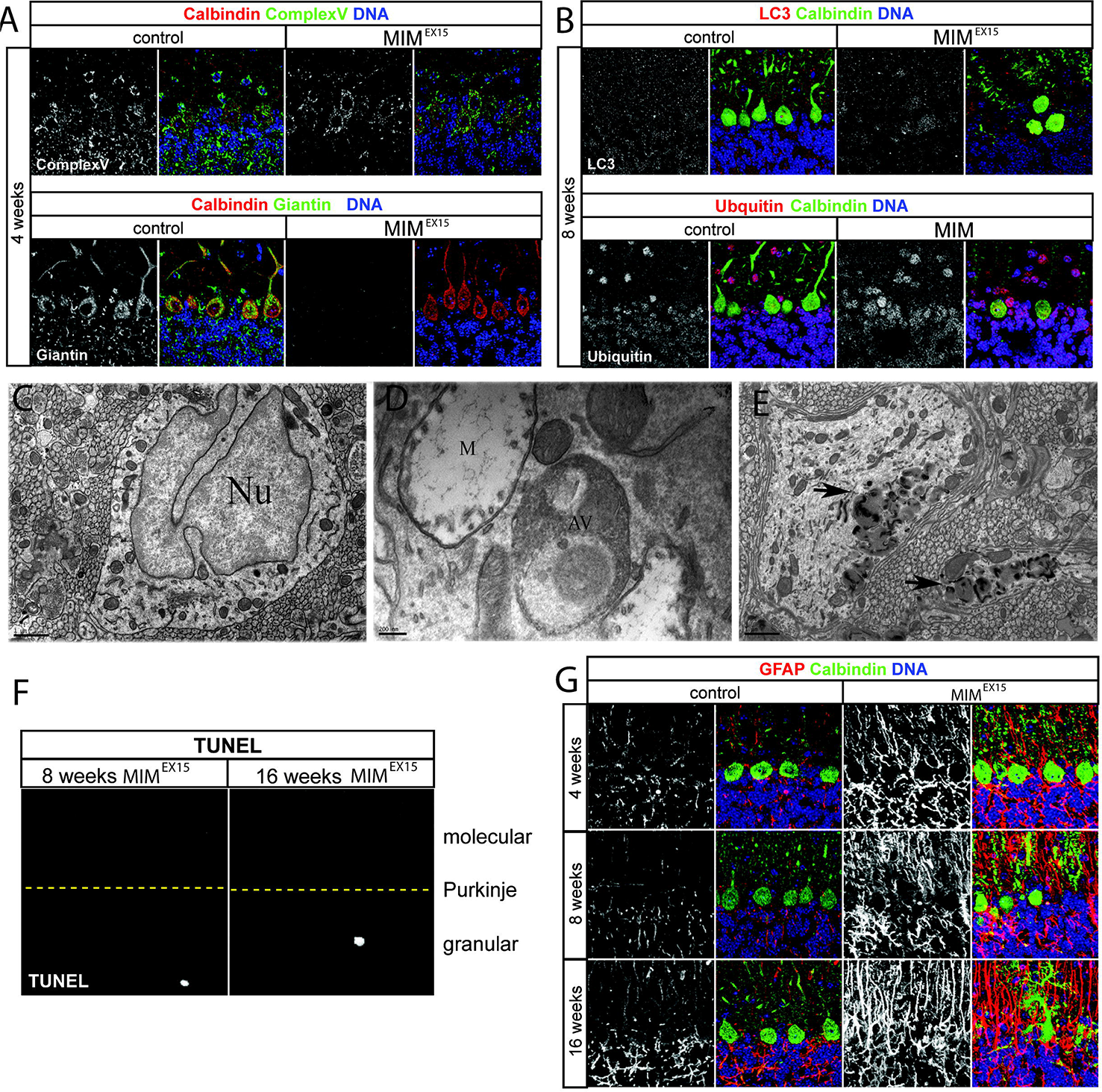
*MIM^EX15^* mutant Purkinje neurons undergo autophagy. **A:** *MIM^EX15^* mutants display fused mitochondria shown by increased Complex 5 ATP-synthase immuno-staining and collapsed Golgi shown by reduced Giantin immune-staining at 4 weeks. **B:** *MIM^EX15^* mutants show increased expression of the autophagocytic markers LC3 and ubiquitin at 8 weeks. **C:** Electron microcopy on 8-week *MIM^EX15^* mutants showed notched Purkinje neuron nuclei (N) (1um scale bar), **D:** swollen autophagocytic mitochondria (M) and autophagic vacuoles (AV) (200nm scale bar). **E:** Lower magnification example of autophagic vacuoles (arrow) in Purkinje neurons (1μm scale bar). **F:** MIM^EX15^ mutant cerebella do not have increased TUNEL stain at 8 or 16 weeks of age. Yellow dotted line denotes Purkinje layer. **G:** MIM^EX15^ mutants show increased GFAP+ glial infiltration during disease progression.

### Mtss1 prevents SFK dependent Purkinje neuron firing defects and ataxia

To characterize cellular changes associated with the ataxia present in 4-week old MIM^EX15^ mice, we examined the dendritic tree of individual biocytin injected Purkinje neurons (**Fig 3A**). Purkinje neuron dendritic arbor collapse has been observed in several SCA models including SCA1 (2), SCA5 (3), while many other models have shown thinned molecular layer including SCA2(1), SCA3 (50), that likely reflects reduced Purkinje dendritic volume. Similarly, *MIM^EX15^* mutants showed a 60% reduction in the expansiveness of the dendritic tree (**Fig 3B**) and a significant decrease in the number of dendritic spines (**Fig 3C**), although no significant difference was detected in spine length (**Fig 3D**) or width (**Fig 3E**).

**Figure 3.**
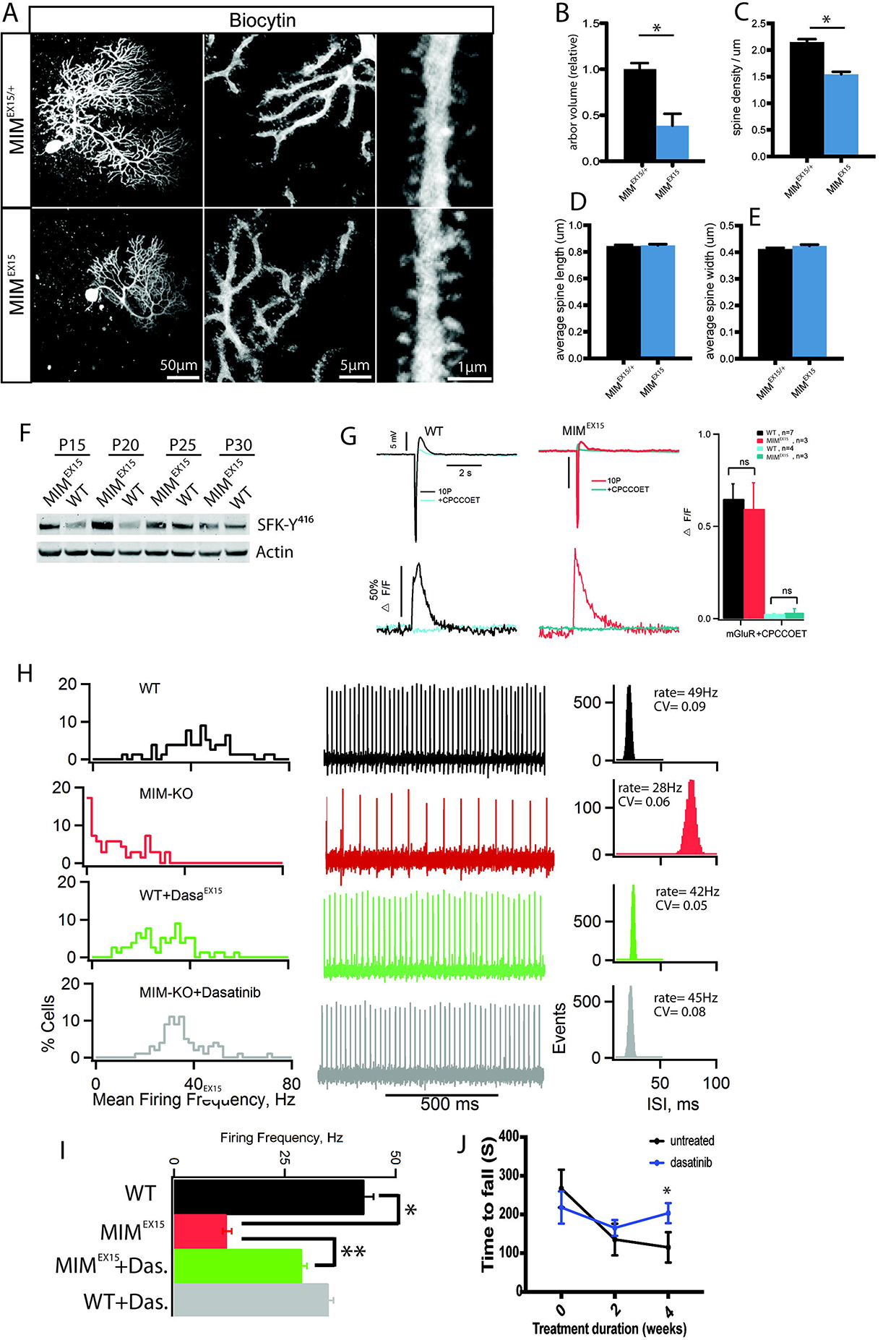
Mtss1 prevents SFK dependent firing defects and ataxia. **A:** Confocal projection of an individual Purkinje cell filled with biocytin and with fluorescent dye to visualize morphology (50μm, 5μm, 1μm scale bars). **B:** Measurement of dye filled Purkinje neurons show *MIM^EX15^* mutants have reduced arbor volume (n=3 each genotype), **C:** reduced dendritic spine density, but D: no change in dendritic spine length and E: no change in dendritic spine width (*MIM^EX15/^+* n=3, 1720 spines; *MIM^EX15^* n=3, 1454 spines, *p<0.05 student’s t-test). Error bars, s.e.m F: Western blot for active SFK-Y416 phosphorylation with Actin loading control. Cerebellar lysate from *MIM^EX15^* and age matched controls collected at indicated times between post-natal day 15 (P15) and post-natal day 30 (P30). G: Slow excitatory post synaptic potential (EPSP) spikes in wild type (WT) and *MIM^EX15^* (top) elicited by stimulation of Parallel Fibers with 10 pulse trains at 100 Hz in the presence AMPA, NMDA and GABA receptor antagonists (control conditions). Corresponding intra-cellular Ca^2^+ signals (ΔF/F) for responses for WT and *MIM^EX15^* mGluR EPSPs are illustrated. EPSPs and corresponding Ca^2^+ signals are blocked by mGluR1 antagonist CPCCOET (bottom). Summary data of intracellular Ca^2^+ signals (ΔF/F) for responses for WT and MTSS1^EX15^ in control conditions and in presence of CPCCOET are shown (right). H: Percent histograms of Purkinje neuron mean firing frequencies (left), examples of extracellular recording of 1 second duration of a spontaneously spiking Purkinje neuron in respective condition (center), and histograms of inter-spike intervals calculated for the 2 minute recording periods of the same neuron (right) are shown for WT, *MIM^EX15^*, WT+dasatinib, or MIM^EX15^+dasatinib conditions. I: Summary of data presented in H *p=4.95E-29 **p=2.1E-14, student’s t-test J: Direct cerebellar administration of dasatinib maintains rotarod performance, slowing the progressive ataxia in *MIM^EX15^* mice. p<0.006 Error bars, s.e.m.

In dermal fibroblasts and *Drosophila* border cells MTSS1 functions to locally prevent ectopic Src kinase activity and *Mtss1* mutant phenotypes can be rescued by genetically removing Src kinase (28, 29). To determine if *Mtss1* acts similarly in Purkinje neurons we evaluated SFK activity levels in cerebellar lysates from *MIM^EX15^* mutants and found elevated levels of SFK^Y416^ (**Fig 3F**) indicative of increased SKF activity. Previous work has shown strong functional interactions between SKF and metabotropic glutamate receptor type I (mGluR1) neurotransmission at parallel fiber synapse (51). To investigate whether MTSS1/SFK modulation of mGluR1 signaling forms the basis of the ataxia, we performed electrophysiological analysis of Purkinje neurons in cerebellar slices from *MIM^EX15^* mice. We evaluated Purkinje neuron response to parallel fiber stimulation using calcium imaging. We found MIM^EX15^ mutant Purkinje neurons responded with a comparable increase of calcium dependent fluorescence to controls, while adding the mGluR1 antagonist CPCCOet abolished these responses (**Fig 3G**). These data support MTSS1 acting post-synaptically to control Purkinje cell function.

Purkinje neurons maintain a cell autonomous tonic firing rate that is essential for their function (52, 53). Since *MIM^EX15^* Purkinje neurons responded normally to parallel fiber stimulation suggesting normal synaptic transmission, we assayed basal firing rate. Purkinje neuron tonic firing rate is highly sensitive to temperature and may vary slightly between investigators (54). In our assays, wild type cells had a mean firing rate of 43±2Hz (n=2 animals, 62 cells), while *MIM^EX15^* mutants exhibited a 12±1Hz mean rate (n=2 animals, 55 cells) (**Figs 3H, 3I**). Previous studies of SCA mouse models demonstrated reduced tonic firing is a basis for ataxia (1, 3, 5). Since basal firing is reduced at an age when *MIM^EX15^* mice possess a normal number of Purkinje neurons, our results suggest neuron malfunction rather than loss underlies the initial ataxia phenotype.

MTSS1/Src double mutants rescue MTSS1 phenotypes in Drosophila and vertebrate cell culture. To test the hypothesis that reducing SFK activity would ameliorate the *MIM^EX15^* ataxia phenotype, we added the FDA-approved SFK inhibitor dasatinib to cerebellar slice preparations and measured basal firing rate, using a concentration approximately 2-fold over *in vivo* IC50 (200nM, Fig 3H, 3I). Dasatinib significantly increased the *MIM^EX15^* basal firing rate from baseline to 29±1Hz (n=2 animals, 62 cells). We also observed that dasatinib slightly reduced the wild type basal firing rate to 35±1Hz (n=2 animals, 79 cells). Time course experiments showed the increase in basal firing rate occurred over 5 hours (**Fig S2**), consistent with a low concentration, high affinity mechanism of action. Direct modulation of ion channel or mGluR1 activity raises basal firing within minutes (4, 55), suggesting that dasatinib works through a distinct mechanism. To determine whether SFK inhibition ameliorates ataxia *in vivo* we administered dasatinib directly to the cerebellum via minipumps to overcome poor CNS bioavailability (56). Over 4 weeks, dasatinib treated *MIM^EX15^* mice were protected from disease progression while untreated mice showed progressively worsening rotarod performance (**Fig 3J**). These results demonstrate that Src family kinases act downstream of MTSS1 and that SFK inhibitors rescue *Mtss1*-dependent basal firing rate defects to slow disease progression.

### *Mtss1* is a translation target of ATXN2

The slow basal firing and ataxia preceding cell death seen in the MIM^EX15^ mutants resembles that seen in other SCA models such as SCA1, SCA2, and SCA5, prompting us to investigate whether MTSS1/SFK dysregulation occurs in other ataxias.SCA2 is caused by an expansion in the polyglutamine (polyQ) tract of the RNA binding protein ATAXIN-2 (ATXN2)(57). The exact molecular defects that drive SCA2 pathogenesis remain unclear, as loss of function mice do not recapitulate the SCA2 phenotype (58), while intermediate expansion alleles are associated with increased risk for frontotemporal dementia (59). Atxn2 has an ancestral role in translation control (7, 60), which may be altered with the SCA2 mutation, but the exact targets have yet to be described.

MTSS1 protein abundance is heavily regulated by metastasis-associated miRs which bind to the MTSS1 3’ untranslated region and reduce steady-state MTSS1 protein levels (61–65) To determine whether MTSS1 protein accumulation is sensitive to Atxn2 we examined the *ATXN2^Q127^mouse* model of SCA2 (1). We found reduced Mtss1 abundance (**Fig 4A, S3**) and increased cerebellar SFK activity in *ATXN2^Q127^* animals compared to wild type littermates (**Fig 4B**).

**Figure 4.**
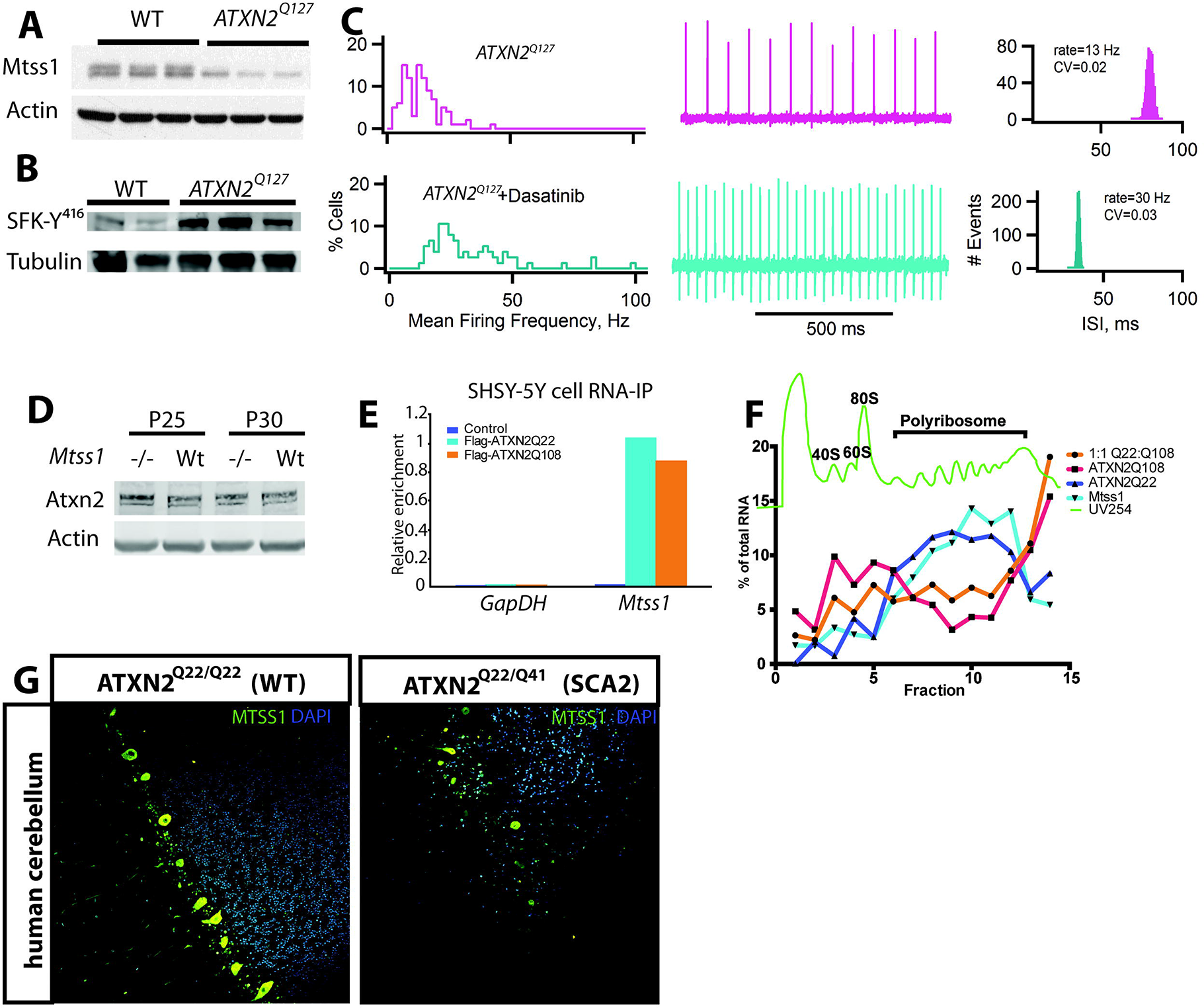
MTSS1 is an Atxn2 translation target. **A:** Western blot of 24-week whole cerebellum lysate shows 90% reduction of upper band that corresponds MTSS1 in ATXN2^Q127^ mice. Actin is included as a loading control **B:** Western blot for active SFK-Y416 phosphorylation with Tubulin loading control using cerebellar lysate from 24 week Atxn2^Q127^ mice **C:** Percent histograms of Purkinje neuron mean firing frequencies (left), examples of extracellular recording of 1 second duration of a spontaneously spiking Purkinje neuron in respective condition (center), and histograms of inter-spike intervals calculated for the 2 minute recording periods of the same neuron for *ATXN2^Q127^* and ATXN2^Q127^+dasatinib ***p=2.355E-14, student’s t-test **D:** Western blot for Atxn2 with tubulin loading control. Cerebellar lysate from post-natal day 25 and 30 *MIM^EX15^* cerebellum and age matched controls. **E:** RNA-IP in SH-SY5Y cells for ATXN2^Q22^ and ATXN2^Q108^ show enrichment for MTSS1 but not GAPDH mRNA **F:** Polyribosome fractionation in 293T cells transfected with MTSS1-UTR reporter and PCDNA, ATXN2^Q22^, ATXN2^Q108^, or ATXN2^Q22^+ATXN2^Q108^. Green line indicates UV254nm absorbance (nucleic acids) with 40S, 60S, 80S, polyribosome peaks labeled. **G:** MTSS1 protein abundance is reduced in human SCA2 cerebellum (Atxn2^Q22/Q41^) Purkinje neurons compared to age matched control (Atxn2^Q22/Q22^).

We sought to determine whether the age-dependent reduction in Purkinje neuron basal firing frequency seen in *ATXN2^Q127^mice* is due to elevated SFK activity. Remarkably, addition of the Src inhibitor to *ATXN2^Q127^* cerebellar slices restored the basal firing rate from an average of 14±1Hz (n=2 animals, 100 cells) to nearly normal levels of 32±2Hz (n=2 animals, 72 cells; **Fig 4C**). As in the MIM^EX15^ mutants, the firing rate reached maximal effect at 5-6 hours of Src inhibition (**Fig S2**), leading us to conclude that inappropriate SFK activity underlies both the ATXN2 and MTSS1-mediated firing phenotype.

The convergence of *Mtss1* and ATXN2 on Src activity suggested they work in a common or parallel molecular pathway. To distinguish between these possibilities, we further interrogated MTSS1 protein levels in *ATXN2^Q127^* cerebella. We found a progressive reduction of MTSS1 protein in ATXN2^Q127^ Purkinje neurons (**Fig 4D**), leading to a 90% reduction at 24 weeks of age (**Fig 4B**, upper band). We failed to see comparable changes in ATXN2 levels in MIMEX15 mice (**Fig 4D**). Because ATXN2 possesses RNA binding activity, and *Mtss1* contains a long 3’UTR, we hypothesized that ATXN2 controls *Mtss1* translation in Purkinje neurons. RNA-IP followed by QPCR showed both WT ATXN^Q22^ and ATXN2^Q108^ specifically bound *MTSS1* mRNA compared to *GAPDH* control. (**Fig 4E**). Using a luciferase reported fused to the MTSS1 3’ UTR we were able to map the ATXN2 interacting domain to a central 500bp region that was sufficient for both RNA-protein interaction and translation control (**Fig S3**). Furthermore, polyribosome fractionation experiments revealed that pathogenic ATXN2^Q108^ was sufficient to block the translation of reporter mRNA fused to the *MTSS1* 3’UTR shifting the transcript from the polyribosome fractions to a detergent resistant fraction consistent with stress granules (**Fig 4F**). These results suggest the pathogenic ATXN2 acts directly as a dominant negative RNA binding protein preventing MTSS1 translation. Notably, we observed MTSS1 abundance is reduced in human SCA patient cerebellum, bolstering the evolutionary conservation of the ATXN2/MTSS1 interaction (**Fig 4G**).

### SFK inhibition rescues Purkinje neuron firing across SCA

Two other SCA mouse models have been shown to have slow basal firing rates, SCA1 (2) and SCA5 (3). Much like SCA2, SCA1 is due to a polyQ expansion in the RNA binding protein ATAXIN-1 (ATXN1)(66). One observed result of the SCA1 allele is changed ATXN1 association with transcriptional regulatory complexes (67), leading to vastly different Purkinje neuron mRNA profiles (68). However, the exact targets that drive SCA1 pathogenesis are still being determined. Unlike SCA1 and SCA2, SCA5 a pure cerebellar ataxia due to lesions in the structural protein β-III spectrin (*SPTNB2*) (13). SPTNB2 directly binds and controls the cell membrane localization of EAAT4 glutamate reuptake receptor which helps clear synaptic glutamate in several types of neurons(12, 69).

If SCA1 or SCA5 acts similarly to SCA2 by regulating the MTSS1/SFK cassette, we would expect decreased MTSS1 abundance and increased active SFK^Y416^. Indeed, in the *ATXN1^Q82^* mouse model of SCA1 (70) we observed a dramatic decrease in MTSS1 protein abundance and corresponding increase in SFK^Y416^ at 15 weeks (**Fig 5A**). Atxn1 pathogenicity is partially driven by phosphorylation at serine 776 (67), which was unchanged in *MIM^EX15^* mice, suggesting MTSS1 is a target of the SCA1 allele (**Fig 5B**). Additionally, *Mtss1* transcript abundance is reduced at multiple ages in *ATXN1^Q8^* mice (68) (**Fig 5C**). We found treating *ATXN1^Q82^* slices with dasatinib increased the basal firing rate from a baseline of 15±1Hz (n=3 animals, 21 cells) to 23±2Hz (n=3 animals, 21 cells), a level statistically indistinguishable from dasatinib-treated controls (**Fig 5C**).

**Figure 5.**
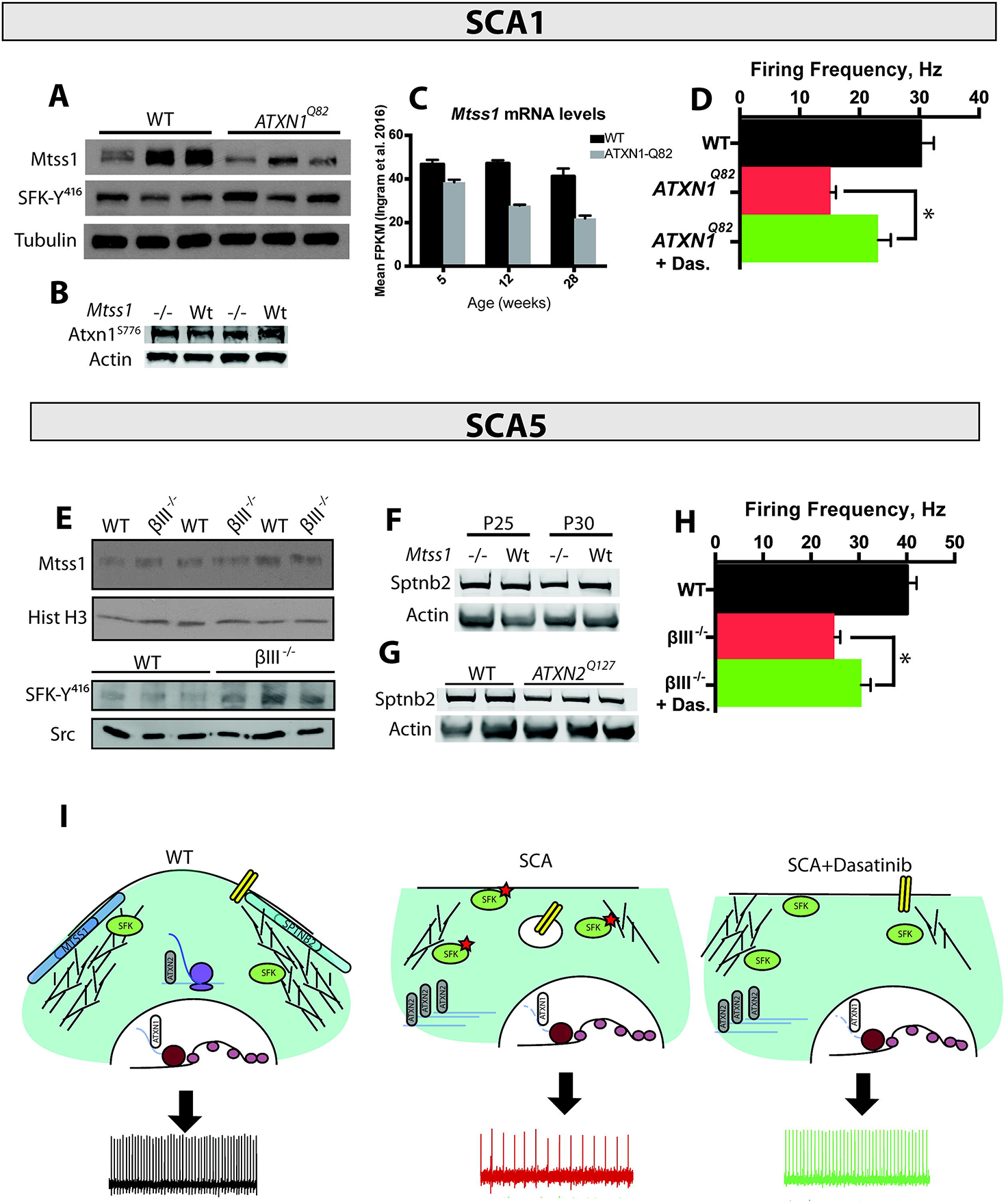
SFK dysregulation occurs in multiple SCA. **A:** Western blot of 15-week whole cerebellum lysate shows reduction of upper band that corresponds MTSS1 in *ATXN1^Q82^* mice, and increase in active Src-Y416 phosphorylation. Tubulin is included as a loading control. **B:** Western blot of 1 month MIM^EX15^ cerebellum lysate shows no change in phospho-Serin776 ATXN1 levels. **C:** RNA-seq from *ATXN1^Q82^* cerebellum shows consistent reduction in FPKM for *Mtss1* mRNA. **D:** Mean firing frequency values in Hz for WT and *ATXN1^Q82^* mice, with and without dasatinib treatment. Error bars, s.e.m. (*p<0.002, student’s t-test) D: Western blot of 15-week whole cerebellum lysate shows no change MTSS1 in βIII-spectrin^-/-^ mice, yet active SFK-Y416 phosphorylation is increased. Histone H3 and total Src are included as a loading controls. E:Western blot in NNN week SPTBN2 null mice show no change in MTSS1 abundance and increased active SFK. F: SPTNB2 abundance is not changed in 1 month MIM^EX15^ mice. G: SPTNB2 levels are not changed in 24 week ATXN2^Q127^ mice. H: Mean firing frequency values in Hz for WT and βIII-spectrin^-/-^ mice, with and without dasatinib treatment. Error bars, s.e.m. (*p<0.05, student’s t-test) I: A model where pathogenic alleles of ATNX1 (ATXN1^Q82^) and ATXN2 (ATXN2^Q42^) prevent the accumulation of MTSS1 and SPTBN2 which restrain SFK activity to prevent neurodegeneration.

By contrast, the *Sptbn2* knockout model of SCA5 (βIII^-/-^)(3),, showed no change in MTSS1 protein abundance yet demonstrated a clear increase in SFK^Y416^ phosphorylation (**Fig 5D**). We fail to see changes in SPTNB2 abundance in *MIM^EX15^* mice, and detect only a subtle decrease in SPTNB2 levels in 24-week *ATXN2^Q127^* mice consistent with reduced Purkinje neuron dendritic arbor size (**Fig 5F, 5G**). These data suggest that SPTNB2 works through a different mechanism than MTSS1, ATXN1, or ATXN2, yet we still observed increased basal firing from 25±1Hz (n=2 animals, 31 cells) to 30±2Hz (n=3 animals, 43 cells) over a 7-hour period of dasatinib treatment (**Fig 5H**).

## Discussion

While SCA gene functions appear heterogeneous, our study establishes a genetic framework to understand how several SCA loci regulate SFK activity to ensure neuronal homeostasis and survival. We identify SPTNB2 and MTSS1, proteins that link the cell membrane and actin cytoskeleton, as negative regulators of Src family kinases.

We show that MTSS1 is a target of the SCA genes ATXN1 and ATXN2 (**Fig 5 I**), and that increased SFK activity from lesions in MTSS1, SPTNB2 (SCA5), ATXN1, and ATXN2 reduces Purkinje neuron basal firing, an endophenotype that underlies multiple ataxias, providing support for the clinical use of SFK inhibitors in many SCA patients.

Our results reveal a central role for the MTSS1/SFK regulatory cassette in controlling neuronal homeostasis and survival. MTSS1 regulation of SFKs has been demonstrated in several migratory cell types including metastatic breast cancer and *drosophila* border cells. This is the first demonstration of the cassette functioning in non-migratory post-mitotic cells. MTSS1 integrates the cell membrane and cytoskeletal response to local signals by serving as a docking site for the kinases and phosphatases that control actin polymerization (71), a process essential for dendritic spine assembly, maintenance and function. In fly border cells, MTSS1-regulated SFK activity polarizes the membrane to spatially detect guidance cues. Similarly, MTSS1 functions in neurons to promote dendritic arborization and spine formation, structures that were shown to be essential for maintaining basal firing frequencies by electrically isolating increasing areas of Purkinje neuron dendrites (54). While a previous study of a MTSS1 hypomorph also detected mild dendritic abnormalities in hippocampal neurons (41), the generation of a true null allele provides a dramatic demonstration of neuronal requirements for MTSS1 function. Other members of the I-BAR family of membrane/cytoskeletal signaling proteins have been implicated in human neurological disorders such as microcephaly (72), but it remains to be determined how they interact with MTSS1.

Disruption of post-transcriptional gene regulation leading to altered proteostasis has recently emerged as a key contributor to neurodegeneration. In the cerebellum, reducing the abundance of the RNA-binding protein Pumilio leads to SCA1-like neurodegeneration through a specific increase in ATXN1 protein levels (73, 74). Yet Pumillio binds hundreds of transcripts to control protein levels (75, 76), suggesting that changing protein abundance of a few key effector genes post-transcriptionally leads to disease. Our data demonstrate that *MTSS1* is a key effector gene whose activity is tightly regulated to prevent Purkinje neuron malfunction. Post-transcriptional control of MTSS1 is disrupted in many disease states such as cancer, where MTSS1 levels are reduced by locus deletion or miRNA overexpression and are associated with increased metastasis and poorer prognosis (62, 77). In Purkinje neurons, the SCA1 ATXN1^Q82^ allele reduces MTSS1 transcript levels. ATXN1 is thought to act as a transcriptional regulator by associating with the transcriptional repressor *Capicua* (CIC) (67), though it remains to be shown whether the ATXN1/CIC complex occupies the *MTSS1* promoter. By contrast, the SCA2 allele ATXN2^Q58^ binds the MTSS1 3’ UTR to prevent ribosome binding and MTSS1 translation, ultimately leading to increased SFK activity. ATXN2 (and the redundant gene ATXN2L) have recently been identified in a large complex of 3’ UTR binding proteins that regulate networks of genes controlling epithelial differentiation and homeostasis (78). Our results suggest other ataxia disease genes that control proteostasis may also regulate MTSS1 abundance, and the strong role for miRNAs controlling MTSS1 abundance in cancer suggest they may also function as effectors of as yet undescribed ataxia loci.

The identification of the SFK suppressor MTSS1 as a novel ataxia locus further reinforces the pathological consequences associated with inappropriate SFK activation in response to a variety of cellular stresses. While the cytoskeletal regulator MTSS1 is an evolutionarily-conserved SFK inhibitor, SFK effects on Purkinje neuron basal firing may derive from the fundamental roles SFKs play in cell homeostasis outside cytoskeletal control. For example, SFK control of translation is implicated in Alzheimer disease, as reducing SFK activity proves beneficial for Alzheimer disease progression (24) due to SFK control of pathogenic Aβ translation (79). SFK impairment of autophagy is seen in models of Amylotrophic lateral schlerosis and Duchenne muscular dystrophy (23, 80). Additionally, reduction of Src kinase expression was identified as a suppressor of SCA1 toxicity in Drosophila ommatidia (81), supporting the need for moderating SFK activity. The pleiotropic effects of inappropriate SFK activity suggests that SFK inhibition may be a critical therapeutic node to slow the progression of multiple neurodegenerative disorders including SCAs. Our work points out the need for future development of neuro-active SFK inhibitor variants, as currently approved Src inhibitors were designed for oncology targets and lack potent central nervous system activity. Further, while we provide data for kinase inhibition to suppress MTSS1 loss, we have previously shown that SFK regulation by regulatory receptor tyrosine phosphatases, or deletion of endocytic adapter proteins can also revert the effects of MTSS1 loss. Given the challenge of developing specific kinase inhibitors, our work opens additional therapeutic classes to alleviate the progression of neurodegenerative diseases.

In summary, the identification of *Mtss1* as a novel recessive ataxia locus extends the physiologic functions requiring the MTSS1/SFK signaling cassette, which include cell polarity, migration, and cancer metastasis. Each of these disparate processes highlight the common role MTSS1 plays integrating the cell membrane and cytoskeletal response to local signals, as the dendritic spine defects seen in MIM^EX15^-mutant Purkinje neurons (**Fig 2A-E**) recalls the loss of directional cell extensions in migrating *Drosophila* border cells (29). They also reinforce the critical need to suppress inappropriate SFK activity, and provide a therapeutic opportunity for otherwise devastating and debilitating diseases.

## Author Contributions

AEO and SXA conceived the project. ASB, SXA, BA performed and interpreted most experiments. PM performed and interpreted all electrophysiology in Mtss1^EX15^ and Atxn2^Q127^ mice. EP and MJ performed and interpreted all electrophysiology and western blots in βIII-spectrin^-/-^ mice. RC and VS performed and interpreted all electrophysiology and western blots in Atxn1^Q82^ mice. SP and DS performed and interpreted MTSS1 western blot and QPCR in ATNX2^Q127^ mice and SH-SY5Y cell RNAIP. SP performed and interpreted MTSS1 staining in human samples. ET quantified biocytin-filled Purkinje data. TSO and SMP contributed ideas and interpreted results. ASB and AEO wrote the manuscript with input from all authors.

## Acknowledgements

The authors would like to acknowledge JoAnn Buchanan for assistance with TEM microscopy, Hak Kyun Kim, Miguel Mata, Peter Sarnow for assistance with ribosome profiling, and SBNFL for assistance with behavior assays. Funding: Oro R01 AR052785, Brown F32 GM105227, Otis R01 NS090930, Jackson The Wellcome Trust 093077

## Supplementary Figure Legends

**Figure S1.**
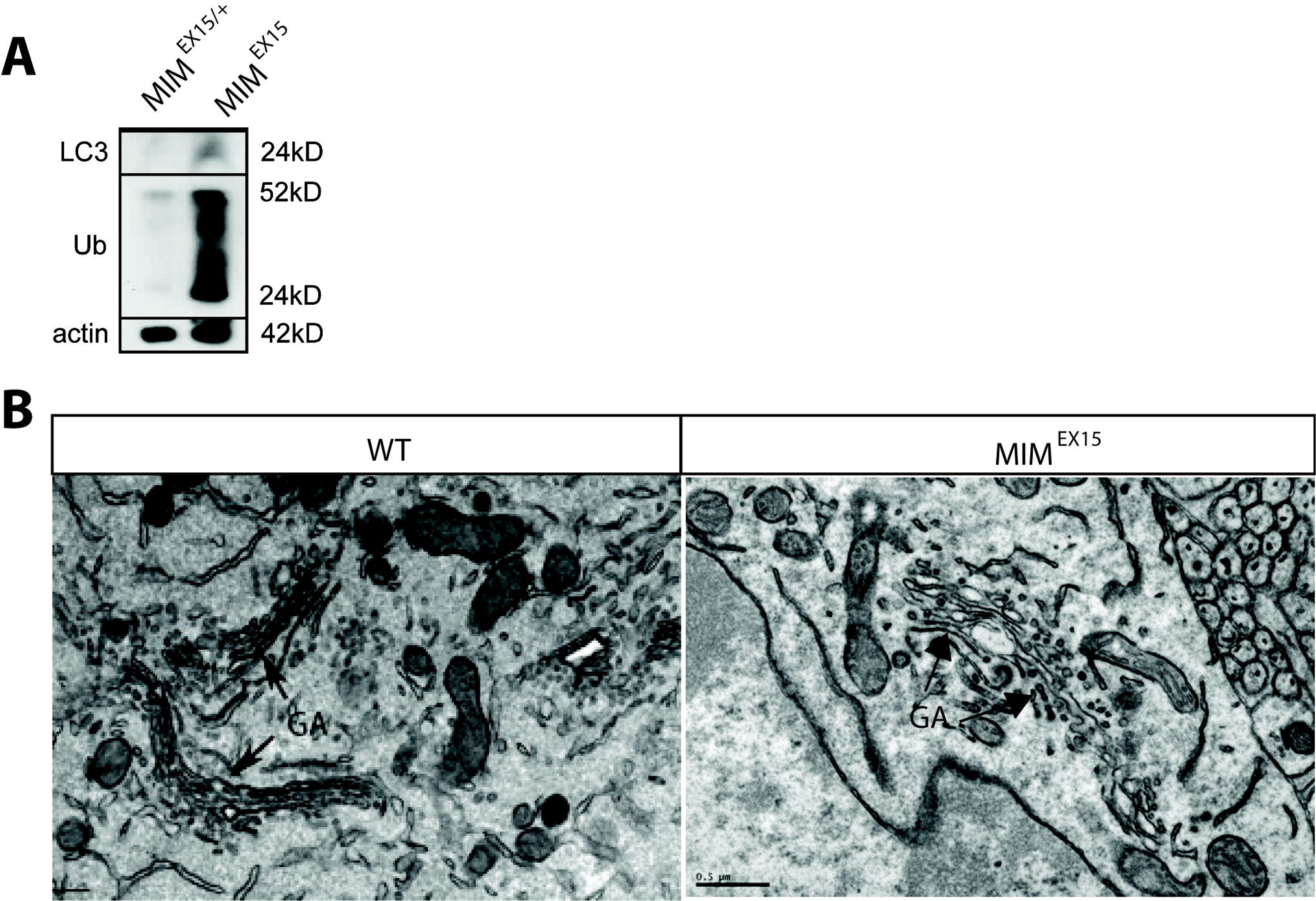
associated with Figure 2 Purkinje cell autophagy in MTSS1 mutants. **A:** Western blot of 8 week cerebellum shows increased LC3 and ubiquitinated protein levels. **B:** Electron micrographs of 8 week cerebellum shows fragmented golgi complex in MIM^EX15^ mutants compared to wild-type.

**Figure S2.**
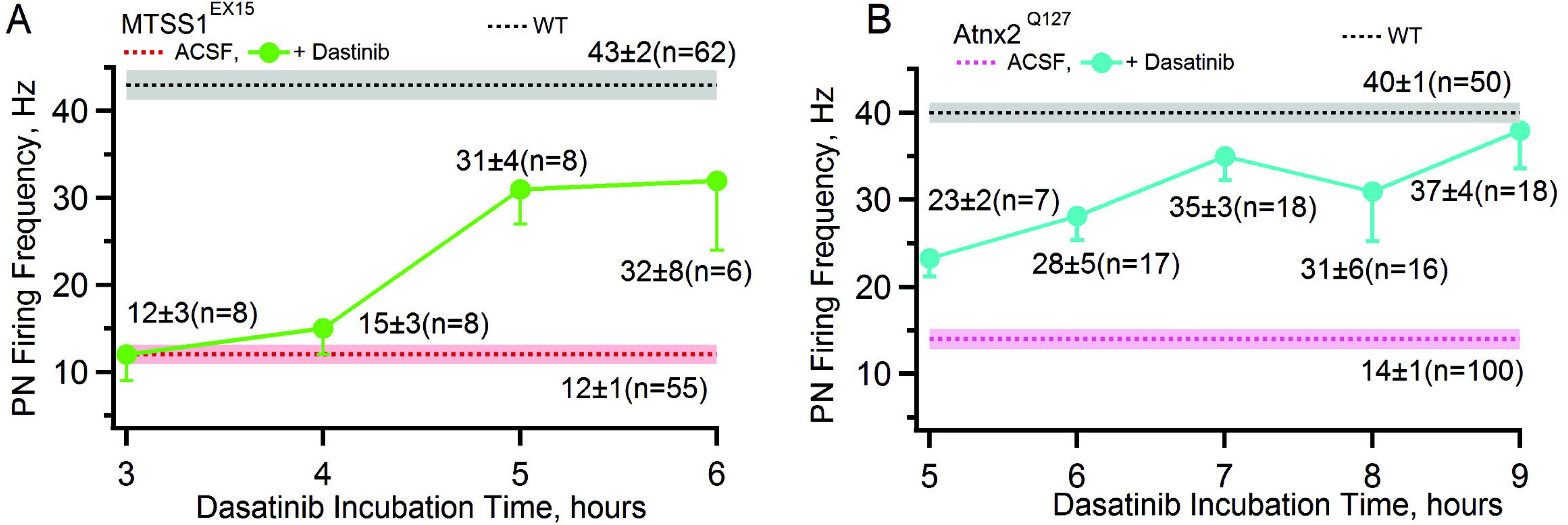
associated with Figures 3 and 4. Acute Src inhibition restores tonic firing rates. Average firing frequency measured at the end of each incubation hour of Dasatinib in MIM^EX15^ **(A)** and ATXN2^Q127^ **(B)** is plotted. A respective mean value with number of Purkinje neurons (PN) at indicated incubations is given in the figure. Note that baseline firing frequency of ATXN2^Q127^ (14±1 Hz, n=100 PNs) mice of this age is similar to the MIM^EX15^ (12±1 Hz, n=55 PNs). Basal firing frequency of PNs from wild type mouse are given.

**Figure S3.**
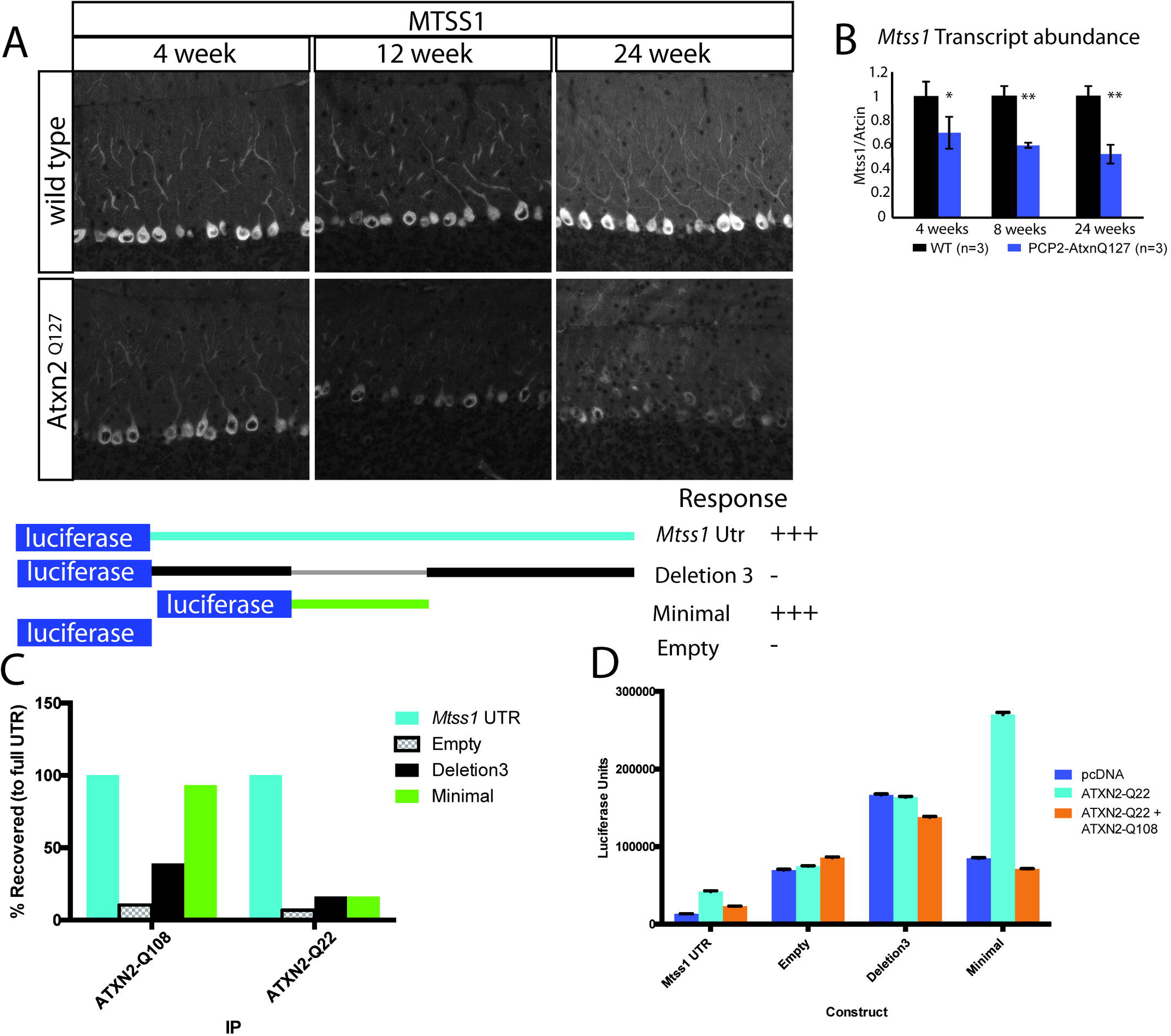
associated with Figure 4. ATXN2 regulates translation through the MTSS1 3’UTR. **A:** MTSS1 immuno-fluorescence in ATXN2^Q127^ and age matched control mice show reduced Purkinje neuron signal at 4, 12 and 24 week time points. **B:** Quantitative RT-PCR (QPCR) shows *Mtss1* transcript abundance is reduced in ATXN2^Q127^ compared to age matched controls. **C:** RNA-IP followed by QPCR show ATXN2 binds the MTSS1 3’UTR. **D:** Luciferase reporter assay show ATXN2 translation is strongly mediated by the 3’UTR, and SCA2 allele is sufficient to block activation.

## EXPERIMENTAL MODEL AND SUBJECT DETAILS

### Generation of MIM ^EX15^ allele

To generate the MIM^EX15^ conditional allele exon15 was cloned into the PGK-gb2 targeting vector between the 5’ LoxP site and the 3’ LoxP/FRT flanking neomycin cassette. The targeting vector contained a 5.97kb 5’ homology arm that included exons 12, 13, 14 and a 2.34kb 3’ homology arm that included the 3’-UTR. The targeting vector was electroporated into C57bl6xSV129 embryonic stem cells, and Neo-resistant colonies were screened by PCR. Chimeric mice were generated by injecting ES cells into blastocysts, and chimeras were mated to a FLP deleter strain(82). To generate MIM^EX15^ null animals, mice with the MIM^EX15^ conditional allele were crossed to HPRT-Cre mice(83). Mice were maintained on a mixed C57bl6 SV129 background and examined at listed ages.

**MIM^CKO^**: MIM ^EX15Loxp^ mice were crossed to L7-Cre (84) to generate MIM^CKO^.

**ATXN2^Q127^**: ATXN2^Q127^ mice were previously characterized in (1).

**ATXN1^Q82^**: ATXN1Q82 mice were previously characterized in (70).

**SPTNB2**: SPTNB2 null mice were previously characterized in (3).

## METHOD DETAILS

### Behavior testing

Rotarod and activity chamber testing was performed by the Stanford Behavioral and Functional Neuroscience Lab. For rotarod mice were trained on 2-20rpm accelerating rod for 4 trials with 15 minute rest intervals between trials. Testing was performed after one rest day at 16rpm constant speed. For activity chamber mice were placed in chamber and measured for 10 minutes 3 times on separate days.

Composite Limb Gait Ledge test was performed as in (43).

### Cerebellar dasatinib administration

Mice were trained on 4-40rpm accelerating rotorod with 15 minute rest intervals. Mice were tested on the same 4-40rpm paradigm after a rest day.

Dasatinib was dissolved in 40% capitsol to a 9mM solution, then diluted in acsf and loaded into azlet pump 1004. Cannulas were inserted at midline, −6.2mm cadual −2.5 DV from bregma. Sutures were closed with ethilon and mice were allowed to recover before subsequent rotarod tests.

### Western blot

Isolated tissues were lysed in RIPA buffer supplemented with complete mini protease inhibitor (Roche) and PhosStop (Roche). Protein concentrations were normalized by using the BCA assay (Pierce). Proteins were electrophoresed on Novex 4-12% or 3-8% gradient gels. Rabbit anti-Src-Y416 (CST 2101S or CST 6943S), mouse anti-beta actin (Sigma), rabbit anti-Sptbn2 (Thermo PA1-46007), rabbit anti-Atxn2 (Sigma HPA021146), mouse anti-Atxn1 (abcam ab63376) primary antibodies were detected with LICOR secondary antibodies.

### Antibodies and Immunofluorescence

Isolated cerebella were immersion fixed in 4% paraformaldehyde and embedded in paraffin. 7μm sections were cut and deparaffinized using standard conditions before staining. Sections were blocked with 20% horse serum 0.3% Triton X-100 in PBS. The following antibodies were used at 1:1000 dilutions:

Rabbit anti-Mtss1(30), Rabbit anti-Calbindin (CST 13176), mouse anti-Calbindin-D-28K monclonal (Sigma), mouse anti-Complex V (Novex 459240), rabbit anti-Ubiquitin (CST 3933), rabbit anti-Giantin (Abcam ab 24586), Chicken anti-GFAP (Abcam ab4674). Alexafluor conjugated secondary antibodies were purchased from Invitrogen. Images were acquired either on a Leica SP2 AOBS laser scanning microscope or a Zeiss axioplan widefield scope.

### Luciferase assay

Luciferase activity was measured using *Mtss1* UTR clone S811096 from Switchgear genomics. Constructs were transfected into 293T cells using FugeneHD 48 hours before measurement. For each well of a 96 well plate, 20ng reporter vector was transfected with reported concentration of ATXN2-Q22-flag or ATXN2-Q108-flag constructs. PCDNA was used to normalize transfection amounts.

Luciferase activity was measured on Molecular Dynamics M5 with 1500ms integration time. Nested deletions were constructed by restriction digest, T4 blunting and ligation. Minimal constructs were generated by Gibson assembly.

### SH-SY5Y RNA-IP

RNA-IP for ATXN2 bound *Mtss1* transcripts in human SH-SY5Y cells was performed as in (7). To identify proteins and RNAs that bind to ATXN2, we carried out protein-RNA immunopre-cipitation (IP) experiments from lysates of SH-SY5Y cells expressing Flag-ATXN2-Q22 and Flag-ATXN2-Q108. Whole cell extracts were prepared by the two-step lyses method. First, cells were lysed with a cytoplasmic extraction buffer (25 mM Tris-HCl pH 7.6, 10 mM NaCl, 0.5% NP40, 2 mM EDTA, 2 mM MgCl_2_, protease and RNAse inhibitors) and cytoplasmic extracts were separated by centrifugation at 14,000 RPM for 20 min. Second, the resultant pellets were suspended in nuclear lysis buffer or high salt lyses buffer (25 mM Tris-HCl, pH 7.6, 500 mM NaCl, 0.5% Nonidet P-40, 2 mM EDTA, 2 mM MgCl2, protease and RNAse inhibitors), and the nuclear extracts were separated by centrifugation at 14,000 RPM for 20 min. The nuclear extracts were then combined with the cytoplasmic extracts and denoted as whole cell extracts. Specifically, while combining cytoplasmic and nuclear extracts, the NaCl concentration was adjusted to physiologic buffer conditions (~150 mM) to preserve in vivo interactions. Ninety percent of cell extracts were subjected to Flag monoclonal antibody (mAb) IP (Anti-Flag M2 Affinity Gel, Sigma Inc.; cat# A2220-1ML) to immunoprecipitate ATXN2 interacting protein-RNA complexes. The remaining 10% of whole cell extracts were saved as the input control for Western blotting and RT-PCR analyses. The IPs were washed with a buffer containing 200 mM NaCl and the bound protein-RNA complexes were eluted from the beads with Flag peptide competition (100 μg/ml).

### MIM 3’UTR RNA-IP

293T Cells were lysed in 20mM Tris Ph 7.5, 140mM NaCl, 1mM EDTA, 10% glycerol, 1% Triton X-100, 20mM DTT supplemented with 20U/ml Superase inhibitor (Life) and complete mini protease inhibitor tablets (Roche). Flag constructs were immunoprecipitated using Anti-Flag magnetic or agarose beads (Sigma). RNA was isolated using Trizol (Life) and treated with DNAse1 (Roche) for 30 minutes. QRTPCR was performed with Brilliant II Sybr green mastermix (Agilent genomics) using the following primers for Luciferase:

F: CTC GGT GCA AGC AGA TGA AC

R: GTT CTC CGC ATG TTT CTC GC

### Fractionation

3 10cm plates of 293T cells were transfected with 1ug *Mtss1* UTR Luciferase, 3ug *Atxn2* 48 hours before fractionation. 1 hour before harvest, media was changed on cells. Ribosomes were stalled by treating cells with 100ug/ml cycloheximide 5 minutes before harvest. Cells were scraped into lysis buffer: 200U/ml Superase, 20mM DTT, 1%Triton-X 100, 20mM Tris pH 7.5, 100mM KCl, 5mM MgCl2,100ug/ml cycloheximide, roche complete mini protease inhibitor tablet, pelleted by centrifuging 8000g 5 minutes. Lysate was normalized by UV 254 absorbance and loaded onto 10%-50% linear sucrose gradients. Gradients were centrifuged 2hrs at 35000rpm in a SW41 rotor. 14 fractions were collected from each gradient using a FoxyR1 collector, and UV254 traces were acquired. For RNA isolation, fractions were treated with TrizolLS (Life), followed by DNAse treatment and QRT-PCR.

### Electrophysiology

#### Preparation of Cerebellar Slices (SCA2 and Mtss1)

Acute parasagittal slices of 285μm thickness were prepared from the cerebella of 4-to 8-week-old mutant and control littermates following published methods(1). In brief, brains were removed quickly and immersed in an ice-cold artificial cerebrospinal fluid (ACSF or extracellular) solution consisting of: 119 mM NaCl, 26 mM NaHCO_3_, 11 mM glucose, 2.5 mM KCl, 2.5 mM CaCl_2_, 1.3 mM MgCl_2_ and 1 mM NaH_2_PO_4_, pH 7.4 when gassed with 5% CO_2_ / 95% O_2_. Cerebella were dissected and sectioned using a vibratome (Leica VT-1000). Slices were initially incubated at 35 °C for 35 min, and then at room temperature before recording in the same ACSF. Dasatinib (200nM) was added during cerebellar sectioning and remained on the slices for recording.

#### Recordings (SCA2 and Mtss1)

Non-invasive extracellular recordings were obtained from Purkinje neurons in voltage-clamp mode at 34.5 ± 1°C. The temperature was maintained using a dual channel heater controller (Model TC-344B, Warner Instruments) and slices were constantly perfused with carbogen-bubbled extracellular solution alone or with 200 nM dasatinib. Cells were visualized with an upright Leica microscope using a water-immersion 40× objective. Glass pipettes were pulled with Model P-1000 (Sutter instruments). Pipettes had 1 to 3 MΩ resistance when filled with extracellular solution and were used to record action potential-associated capacitative current transients near Purkinje neuron axon hillock with the pipette potential held at 0 mV. Data was acquired at 20 kHz using a Multiclamp 700B amplifier, Digidata 1440 with pClamp10 (Molecular Devices), filtered at 4 kHz. A total of 50 to 100 Purkinje neurons were measured from each genotype and each recording was of 2 minutes in duration. The experimenter was blinded to the mouse genotype and 2 to 4 mice were used per genotype. Simultaneous mGluR EPSPs and calcium were measured in the presence of GABA_A_ receptor antagonist, picrotoxin (PTX at 100 μM), AMPA receptor blockers (5 μM NBQX and 10 μM DNQX) using a two-photon microscope and a standard electrophysiology set-up. The patch pipettes had 4 to 5 MΩ resistance when filled with internal solution (135 mM KMS0_4_, NaCl, 10 mM HEPES, 3 mM MgATP, 0.3 mM Na2GTP) containing 200 μM Oregon Green Bapta1 and 20 μM Alexa 594. The stimulating electrode was filled with ACSF containing 20 μM Alexa 594, placed in the dendritic region to minimally stimulate PF synaptic inputs. Slow mGluR EPSPs in control littermate and mutant were elicited by stimulation of PFs with 100 Hz trains, and 10 pulses in the presence of receptor antagonists that block AMPA, NMDA, GABA_A_ receptors. Corresponding intracellular Ca^2^+ signals (ΔF/F) for responses for wild type and mutant mGluR EPSPs were blocked by the mGluR1 antagonist CPCCOET.

Experiments were analyzed using both the Clampfit and Igor algorithms, and were further analyzed using Microsoft Excel. Figures were made in Igor program. Calcium signals were analyzed using Slidebook (Intelligent Imaging Innovations, Inc.). Results are presented as mean ±SEM. All chemicals were purchased either from Sigma Aldrich, Tocris and Invitrogen, USA.)

### Biocytin fills of Purkinje neurons or Intracellular labeling of Purkinje neurons with Biocytin

Biocytin filling of Purkinje neurons was performed using recording pipettes filled with 1% Biocytin (Tocris). Purkinje neurons were filled for 15 to 30 minutes and then the pipette was removed slowly for enabling the cell membrane to reseal. Slices were then fixed in 4% Paraformaldehyde overnight and washed 3 times with phosphate-buffered saline (PBS). Slices were then incubated with Alexa Fluor 488 streptavidin (1:500, Life S11223) in PBS, 0.5% Triton X-100, and 10% normal goat serum for 90min. After another 3 PBS washes, the slices were then mounted onto a slide with prolong gold. Individual biocytin-filled Purkinje cells were visualized on a Leica SP2 AOBS laser scanning microscope at a 0.5um step size. Dendritic arbor volume was measured by calculating the biocytin-filled area in each confocal optical section using ImageJ, adding the areas in each z-stack, and multiplying by the step size.

### *Ex-vivo* Electrophysiology (SCA1)

#### Solutions

Artificial CSF (aCSF) contained the following (in mM): 125 NaCl, 3.5 KCl, 26 NaHCO_3_, 1.25 NaH_2_PO_4_, 2 CaCl_2_, 1 MgCl_2_, and 10 glucose. For all recordings, pipettes were filled with internal recording solution containing the following (in mM): 119 K Gluconate, 2 Na gluconate, 6 NaCl, 2 MgCl_2_, 0.9 EGTA, 10 HEPES, 14 Tris-phosphocreatine, 4 MgATP, 0.3 tris-GTP, pH 7.3, osmolarity 290 mOsm.

#### Preparation of brain slices for electrophysiological recordings

Mice were anesthetized by isofluorane inhalation, decapitated, and the brains were submerged in pre-warmed (33°C) aCSF. Slices were prepared in aCSF containing dasatinib or DMSO and held at 32.5-34°C on a VT1200 vibratome (Leica). Slices were prepared to a thickness of 300 μm. Once slices were obtained, they were incubated in continuously carbogen (95% 0_2_/5% CO_2_)-bubbled aCSF containing DMSO or dasatinib for 45 minutes at 33°C. Slices were subsequently stored in continuously carbon-bubbled aCSF containing DMSO or dasatinib at room temperature until use. For recordings, slices were placed in a recording chamber and continuously perfused with carbogen-bubbled ACSF containing DMSO or dasatinib at 33°C with a flow rate of 2—3 mls/min.

#### Patch-clamp recordings

Purkinje neurons were identified for patch-clamp recordings in parasagittal cerebellar slices using a 40x water-immersion objective and Eclipse FN1 upright microscope (Nikon) with infrared differential interference contrast (IR-DIC) optics that were visualized using NIS Elements image analysis software (Nikon). Borosilicate glass patch pipettes were pulled with resistances of 3-5 MΩ. Recordings were made 5 hours after slice preparation. Data were acquired using an Axon CV-7B headstage amplifier, Axon Multiclamp 700B amplifier, Digidata 1440A interface, and pClamp-10 software (MDS Analytical Technologies). In all cases, acquired data were digitized at 100 kHz.

## QUANTIFICATION AND STATISTICAL ANALYSIS

For cell counts, firing rates, rotorod and activity chamber 2-tailed non-homodidactic Student’s T-test was used to calculate significance.

For cerebellar dasatinib cohorts of 2-5 mice were tested. Analysis was T-test with two-stage step-up method of Benjamini, Krieger and Yekutieli with a 5% FDR for multiple test correction.

For electrophysiology 2-3 mice per condition were evaluated with investigator blinded to genotype. For MTSS1 rotorod and activity chamber cohorts of 10 age matched animals were examined with investigator blind to genotypes.

For western blots and immune fluorescence 2-3 mice per genotype and age were evaluated.

